# Identification of five antiviral compounds from the Pandemic Response Box targeting SARS-CoV-2

**DOI:** 10.1101/2020.05.17.100404

**Authors:** Melle Holwerda, Philip V’kovski, Manon Wider, Volker Thiel, Ronald Dijkman

## Abstract

With currently over 4 million confirmed cases worldwide, including more than 300’000 deaths, the current Severe Acute Respiratory Syndrome Coronavirus 2 (SARS-CoV-2) pandemic has a major impact on the economy and health care system. Currently, a limited amount of prophylactic or therapeutic intervention options are available against SARS-CoV-2. In this study, we screened 400 compounds from the antimicrobial ‘Pandemic Response Box’ library for inhibiting properties against SARS-CoV-2. We identified sixteen compounds that potently inhibited SARS-CoV-2 replication, of which five compounds displayed equal or even higher antiviral activity compared to Remdesivir. These results show that five compounds should be further investigated for their mode of action, safety and efficacy against SARS-CoV-2.

**Highlights:**

- 400 compounds from the pandemic response box were tested for antiviral activity against SARS-CoV-2.
- 5 compounds had an equal or higher antiviral efficacy towards SARS-CoV-2, compared to the nucleoside analogue Remdesivir.

## Introduction

In December 2019, a new zoonotic coronavirus emerged in Wuhan, Hubei Province, China named Severe Acute Respiratory Syndrome Coronavirus 2 (SARS-CoV-2) which is the etiological agent of Coronavirus Disease 2019 (COVID-19) [1–3]. The clinical features of SARS-CoV-2-infected patients range from mild cold-like symptoms to severe illness ultimately leading to acute respiratory distress syndrome (ARDS) [2,4]. Patients at older age and with underlying comorbidities are at higher risk for developing severe courses of COVID-19 [5]. Despite unprecedented international public health response measures to contain SARS-CoV-2 transmissions, the viral outbreak is currently categorized as a pandemic with over 4 million confirmed-laboratory cases reported worldwide, including over 300,000 deaths as of 8^th^ of May 2020 [6]. At present, and despite earlier outbreaks by SARS-CoV and Middle East Respiratory Syndrome (MERS)-CoV, there are limited approved antiviral treatment options such as antiviral drugs, vaccines and immuno-prophylaxis that can be used prophylactically or therapeutically to halt the current SARS-CoV-2 infections.

Vaccine development takes multiple years until it can be administered to patients, and although these processes are expedited during outbreaks of emerging viruses, the eventual worldwide vaccine distribution may be delayed for several months after the initial outbreak [7]. Moreover, while vaccines are used prophylactically, antiviral drugs can be employed both prophylactically and therapeutically. For SARS-CoV-2, several antiviral compounds are currently evaluated, such as the nucleoside analogue Remdesivir, the TMPRSS2 protease inhibitor cameostat mesylate, and the antimalaria drug (Hydroxy-) chloroquine, that target different stages of the viral replication cycle [8,9]. These three antiviral drugs have been recently tested clinically in small cohorts, and their efficacy against SARS-CoV-2 infections is currently evaluated [10]. Nevertheless, RNA viruses, including coronaviruses, are known to rapidly evade antiviral drug inhibition by developing resistance mutations and subsequent selection of drug-resistant viral populations [11–13]. Therefore, the use of multiple drug regimes as well as expanding the repertoire of available antiviral treatment options are of crucial importance to combat the SARS-CoV-2 pandemic.

Since the beginning of the 21^st^ century, the world has encountered multiple epidemics that were caused by a viral or bacterial agent, like the Ebola-, Measles-, Zika-viruses and cholera [14]. Some epidemics even reached pandemic proportions, like the Influenza A/H1N1 virus and the currently circulating SARS-CoV-2. As a rapid response to these virulent agents, the Medicines for Malaria Venture (MMV, mmv.org) and Drugs for Neglected Diseases intitiative (DNDi, dndi.org) developed the pandemic response box (PRB), a compound library containing 400 compounds with antibacterial, antifungal and antiviral properties. This compound library contains drugs that are already marketed or are at various stages of development and analysis and allows for the rapid investigation of repurposing drugs against newly emerging pathogens.

To this end, we performed an *in vitro*-based screen of 400 preselected compounds with antibacterial, antifungal and antiviral properties contained in the PRB and assessed their antiviral activity against SARS-CoV-2. A stringent large-scale screen in Vero-E6 cells highlighted sixteen compounds that prevented virus-induced cytopathogenic effects (CPE) while displaying low cytotoxicity and no effect on cell viability. Further evaluation revealed that five compounds had an equal or higher antiviral efficacy against SARS-CoV-2, compared to the nucleoside analogue Remdesivir. Together, these data demonstrate that stringent *in vitro* screening of preselected compounds leads to the rapid identification of potent antiviral candidate drugs targeting SARS-CoV-2.

## Material and methods

### Cells, viruses and compounds

Vero-E6 cells (kindly provided by M. Müller/C. Drosten, Charité, Berlin, Germany) were propagated in Dulbecco’s Modified Eagle Medium-GlutaMAX, 10% (v/v) heat-inactivated fetal bovine serum, 100 mg/ml streptomycin, 100 IU/ml penicillin, 1% (w/v) non-essential amino acids and 15 mM HEPES (Gibco). Cells were maintained at 37°C in a humidified incubator with 5% CO_2_. SARS-CoV-2 (SARS-CoV-2/München-1.1/2020/929 [15], stocks were produced on Vero-E6 cells, aliquoted and stored at - 80°C. Viral titers were determined by tissue culture infectious dose 50 (TCID_50_) on Vero-E6 cells.

### Compound preparation of the pandemic response box

All compounds were dissolved and diluted in dimethylsulfoxide (DMSO, Sigma) to 1 mM aliquots in 96-well plates (TPP) that were kept at −20°C until further use. Compounds were diluted at the indicated concentration in cell culture medium.

### Survival screening

Vero-E6 cells were seeded in 96-well clear bottom, black plates (Costar), 20’000 cells per well one day prior to the experiment. Cells were pretreated for 2 hours with 1 µM of each compound contained in the PRB. Remdesivir [8], K22 [12] and DMSO controls were included in each plate. Subsequently, cells were infected with SARS-CoV-2 at a multiplicity of infection (MOI) of 0.01 in compound-containing medium and incubated at 37°C in a humidified incubator with 5% CO_2_. Uninfected (mock) controls were included in each plate. At 48 hours post-infection (hpi), cells were visually inspected for virus-induced CPE prior to fixation with 4% (v/v) neutral buffered formalin (Formafix AG) and stained with crystal violet. Cell viability and cytotoxicity were assessed in parallel, in identically treated, uninfected plates. Two independent experiments were performed, each including a technical duplicate. Wells containing an intact cell layer without apparent CPE after infection and displaying high cell viability and low cytotoxicity were considered as hits.

### Cell cytotoxicity and cell viability

Cell cytotoxicity and viability were assessed using CellTox(tm) green cytotoxicity assay (Promega) and Cell-Titer Glo 2.0 assay (Promega), respectively, according to manufacturer’s protocols. Readout was performed on a Cytation 5 Cell Imaging Multi-Mode Reader (Biotek).

### IC_50_ determination of selected compounds

Vero-E6 cells were seeded in 96-well clear bottom, black plates (Costar), 20’000 cells per well one day prior to the experiment. Cells were pretreated for 2 hours with 2-fold serial dilutions of selected compounds, ranging from 4 µM to 0.063 µM. Cells were infected with SARS-CoV-2 (MOI of 0.01) in compound-containing medium and incubated at 37°C in a humidified incubator with 5% CO_2_. Cells were fixed with 4% (v/v) neutral buffered formalin at 24 hours post-infection and processed for immunofluorescence analysis. Briefly, cells were permeabilized with 0.1% (v/v) Triton X-100 (Merck) for 5 minutes and blocked in PBS supplemented with 50 mM NH_4_Cl, 0.1% (w/v) Saponin (Sigma) and 2% (w/v) Bovine serum albumine (IgG-free, Jackson immunoresearch). SARS-CoV-2 antigen-positive cells were detected using a rabbit polyclonal anti-SARS-CoV nucleocapsid protein (Rockland, 200-401-A50) and a secondary Alexa Fluor® 488-labeled donkey anti-Rabbit IgG (H+L) (Jackson Immunoresearch). Samples were counterstained using 4’,6-diamidino-2-phenylindole (DAPI, Thermo Fisher Scientific) to visualize the nuclei and finally washed with PBS.

Images were acquired on a Cytation5 Cell Imaging Multi-Mode Reader (Biotek) equipped with a 4x air objective (NA: 0.13). Four images per well were acquired to cover the entire surface of the well and processed and stitched using the Gen5 Image prime software package (v3.08.01). Mean intensity ratios of the GFP (SARS-Nucleocapsid) and DAPI (Nuclei) signals were calculated for each individual well. Cell viability and cytotoxicity were assessed in parallel, in identically treated, uninfected plates.

### Data representation

Graphs were generated using GraphPad Prism software version 8.4.2 and the final figures were assembled in Adobe Illustrator CS6. Brightness and contrast of microscopy picture were minimally adjusted and processed identically to their corresponding control using FIJI. Images were assembled using the FigureJ plugin in FIJI [16].

## Results

### Survival screen with compounds included in the pandemic response box against SARS-CoV-2

To identify potential compounds that can be used as intervention option against SARS-CoV-2 replication, we screened 201 antibacterial, 46 antifungal and 153 antiviral molecules included in the PRB for their antiviral activity. We initially screened the 400 compounds at a conservative concentration of 1 µM. Based on the documented inhibition of coronavirus replication, Remdesivir and K22 were included as a positive control [8,12]. Vero-E6 cells were pretreated for 2 hours and subsequently infected with SARS-CoV-2 (MOI of 0.01) for 48 hours in drug-containing medium. Cell survival was arbitrarily scored, from 0 to 2, upon visual inspection and evaluation of SARS-CoV-2-induced CPE (**Supp. Fig. 1a)**. This screen resulted in a total of five compounds that completely inhibited SARS-CoV-2-induced CPE (white), while twelve other compounds showed an intermediate CPE reduction (Grey). In parallel, we performed cell viability and cell cytotoxicity assays to exclude any detrimental effect of each compound on the cells. Only for one compound (Plate B, D10) we observed low cell viability and high cytotoxicity. The sixteen remaining compounds are antifungal (three), antibacterial (six) and antiviral (seven) compounds **(Table 1)**, which, similarly to their vehicle control (DMSO), Remdesivir and K22, did not influence cell cytotoxicity and cell viability (**Supp. Fig. 1b, c**). These results provide evidence for the relevance of a conservative and rapid screening of libraries containing compounds that target viral replication.

**Table 1:** The sixteen compounds that showed inhibition of the SARS-CoV-2 after the survival screen. Remdesivir was included as reference drug control.

| Plate position: | Section | MMV-number | CHEMBL ID | Websitelink or literaturelink: | Name | Clinical stage according ChEMBL | Chemical formula |
| --- | --- | --- | --- | --- | --- | --- | --- |
| Plate A, D2 | Antifungals | MMV1634386 | CHEMBL3311228 | https://www.ebi.ac.uk/chembl/compound_report_card/CHEMBL3311228/ | Oteseconazole | 3 (Phase III) | C23 H16 F7 N5 O2 |
| Plate A, E2 | Antifungals | MMV637528 | CHEMBL64391 | https://www.ebi.ac.uk/chembl/compound_report_card/CHEMBL64391/ | Itraconazole | 4 (approved) | C35 H38 Cl2 N8 O4 |
| Plate B, A2 | Antibacterials | MMV1483032 | CHEMBL243644 | https://www.ebi.ac.uk/chembl/compound_report_card/CHEMBL243644/ | AC1MTT7T | 0 (research) | C22 H18 N4 O4 |
| Plate B, A7 | Antibacterials | MMV637945 | CHEMBL403 | https://www.ebi.ac.uk/chembl/compound_report_card/CHEMBL403/ | Sulbactam | 4 (approved) | C8 H11 N O5 S |
| Plate B, D9 | Antibacterials | MMV1582492 | CHEMBL3109593 | https://www.ebi.ac.uk/chembl/compound_report_card/CHEMBL3109593/ | Retro-2.1 | 0 (research) | C23 H18 F N3 O S2 |
| Plate B, F3 | Antibacterials | MMV1582487 | CHEMBL198796 | https://www.ebi.ac.uk/chembl/compound_report_card/CHEMBL198796/ | Decylphosphinate | 0 (research) | C13 H28 N O4 P |
| Plate C, A5 | Antibacterials | MMV1578576 | CHEMBL1568820 | https://www.ebi.ac.uk/chembl/compound_report_card/CHEMBL1568820/ | - | 0 (research) | C15 H12 F N3 O |
| Plate C, E7 | Antibacterials | MMV000008 | CHEMBL76 | https://www.ebi.ac.uk/chembl/compound_report_card/CHEMBL76/ | Chloroquine | 4 (approved) | C18 H26 Cl N3 |
| Plate D, A11 | Antivirals | MMV1593513 | CHEMBL408500 | https://www.ebi.ac.uk/chembl/compound_report_card/CHEMBL408500/ | N-Nonyl Deoxynojirimycin (NN-DNJ) | 0 (research) | C15 H31 N O4 |
| Plate D, C7 | Antivirals | MMV690621 | - | Patent: WO2006118607A2 | NA for racemic | - | C18 H16 Cl N3 O |
| Plate D, E10 | Antivirals | MMV1634401 | CHEMBL1652119 | https://www.ebi.ac.uk/chembl/compound_report_card/CHEMBL1652119/ | PBDNJ0803 | 0 (research) | C20 H33 N O5 |
| Plate D, E11 | Antivirals | MMV002780 | CHEMBL402487 | https://www.ebi.ac.uk/chembl/compound_report_card/CHEMBL402487/ | Noscapine | 0 (research) | C22 H23 N O7 |
| Plate E, A6 | Antivirals | MMV1580482 | CHEMBL2436978 | https://www.ebi.ac.uk/chembl/compound_report_card/CHEMBL2436978/<br>Patent: WO 2014085795 A1 | URMC-099-C | 0 (research) | C27 H27 N5 |
| Plate E, B7 | Antivirals | MMV1593544 | CHEMBL3752642 | https://www.ebi.ac.uk/chembl/compound_report_card/CHEMBL3752642/ | - | 0 (research) | C36 H43 N3 O5 |
| Plate E, E11 | Antifungals | MMV002350 | CHEMBL561 | https://www.ebi.ac.uk/chembl/compound_report_card/CHEMBL561/ | Lomefloxacin | 4 (approved) | C17H19F2N3O3 |
| Plate E, F11 | Antivirals | MMV1580483 | CHEMBL3960662 | https://www.ebi.ac.uk/chembl/compound_report_card/CHEMBL3960662/ | AZD0156 | 0 (research) | C26 H31 N5 O3 |
| - | Control | - | CHEMBL4065616 | https://www.ebi.ac.uk/chembl/compound_report_card/CHEMBL4065616/ | Remdesivir | 2 (Phase II) | C27 H35 N6 O8 P |

### Antiviral efficacy against SARS-CoV-2

To further confirm and evaluate the extent of antiviral activity of the previously highlighted sixteen compounds, cells were pretreated with the selected compounds at dosages ranging from 4 µM to 0.063 µM and infected with SARS-CoV-2 (MOI of 0.01). After 24 hours of infection, cells were fixed and processed for immunofluorescence analysis using the anti-SARS-CoV nucleocapsid protein antibody and DAPI. The efficacy of the selected compounds to inhibit SARS-CoV-2, as well as their individual effects of cell viability and cytotoxicity, were compared to Remdesivir. The IC_50_ values for each compound were inferred by calculating the ratio of the total intensity of the nucleocapsid protein and DAPI signals (infected cells / total cells). This indicated that five of sixteen selected candidate compounds, *N*-Nonyldeoxynojirimycin (NN-DNJ), PDNJ0803, Chloroquine, Retro-2.1 and URMC-099-C, inhibited the virus at least as potently as the reference compound Remdesivir **(Figure 1)**.

**Figure 1:**
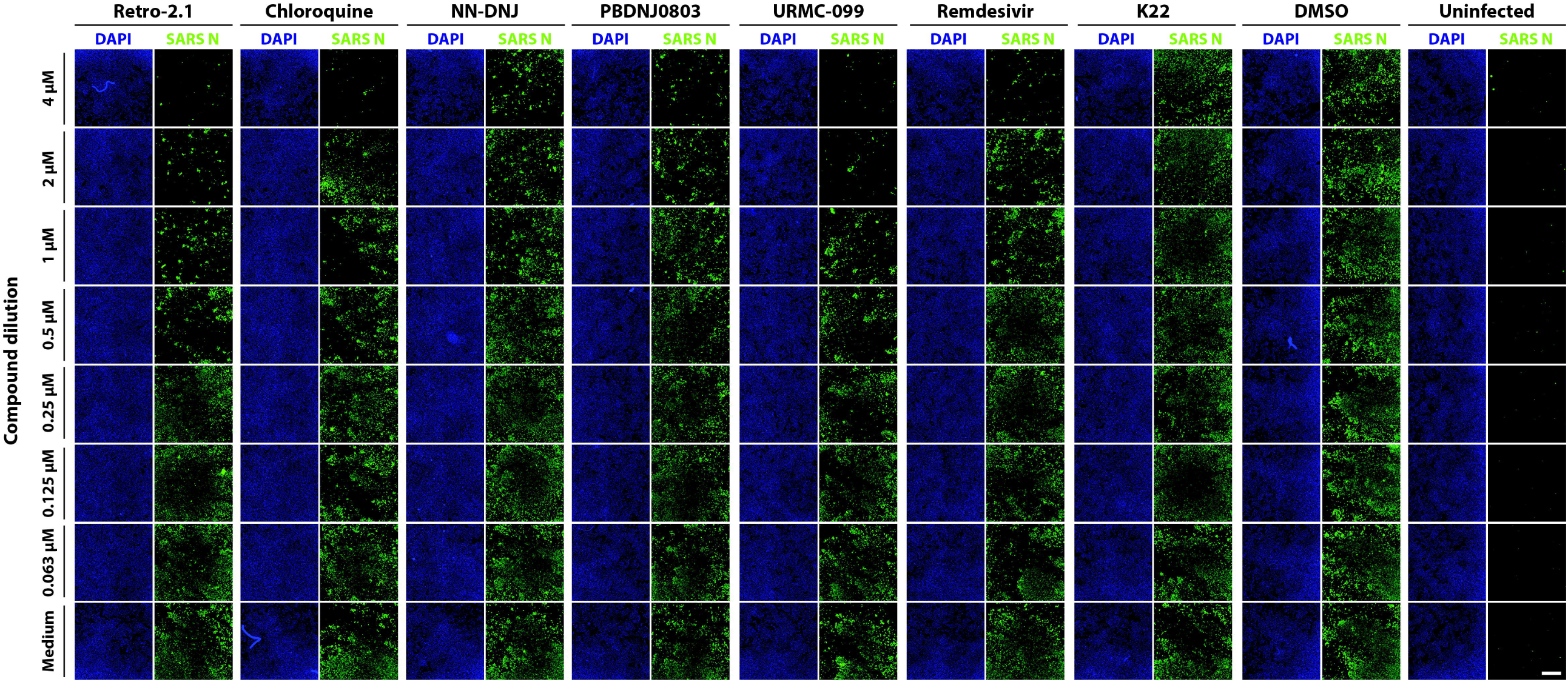
Immunofluorescence staining of the dilution series of the hits. To determine the efficiency of inhibition of each specific compound that were selected after the survival screen, dilution series ranging from 4 to 0.063 µM were prepared followed by infection with SARS-CoV-2 (MOI of 0.01). Vero-E6 cells were pretreated with the compound 2 hours before infection at 37°C with an CO_2_-humidity of 5% and fixed 24 hours post infection, followed by immunostaining with the cross-reactive SARS-CoV Nucleoprotein antigen (SARS-N) and DAPI. Remdesivir and K22 included were included as controls. The acquisition was performed with a 4x air objective whereby four pictures per well were stitched with the build-in Gen5-software. The image are representative of the results of three individual experiments. Scale bar represent 1 μM.

Moreover, these five compounds showed equal or lower IC_50_ values against SARS-CoV-2 replication than the reference compound Remdesivir (IC_50_ = 1.842), without increased cytotoxicity or decreased cell viability compared to the vehicle control on the analyzed concentrations **(Fig. 2 A-G, Table 2)**. The remaining eleven compounds showed little or no inhibition when compared to the reference compound, and therefore no reliable IC_50_ could be calculated **(Supp. Fig 2)**.

**Table 2:**
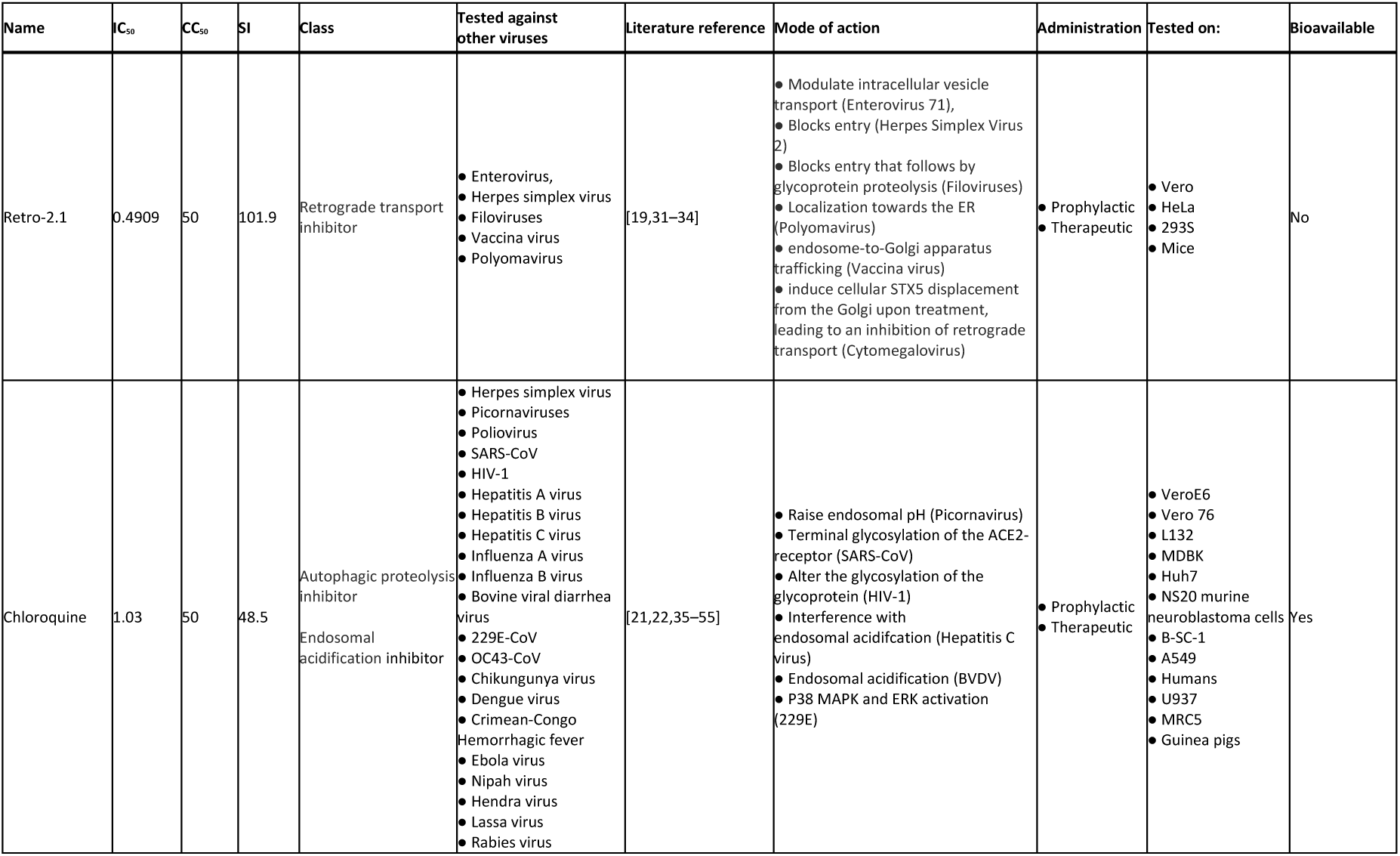
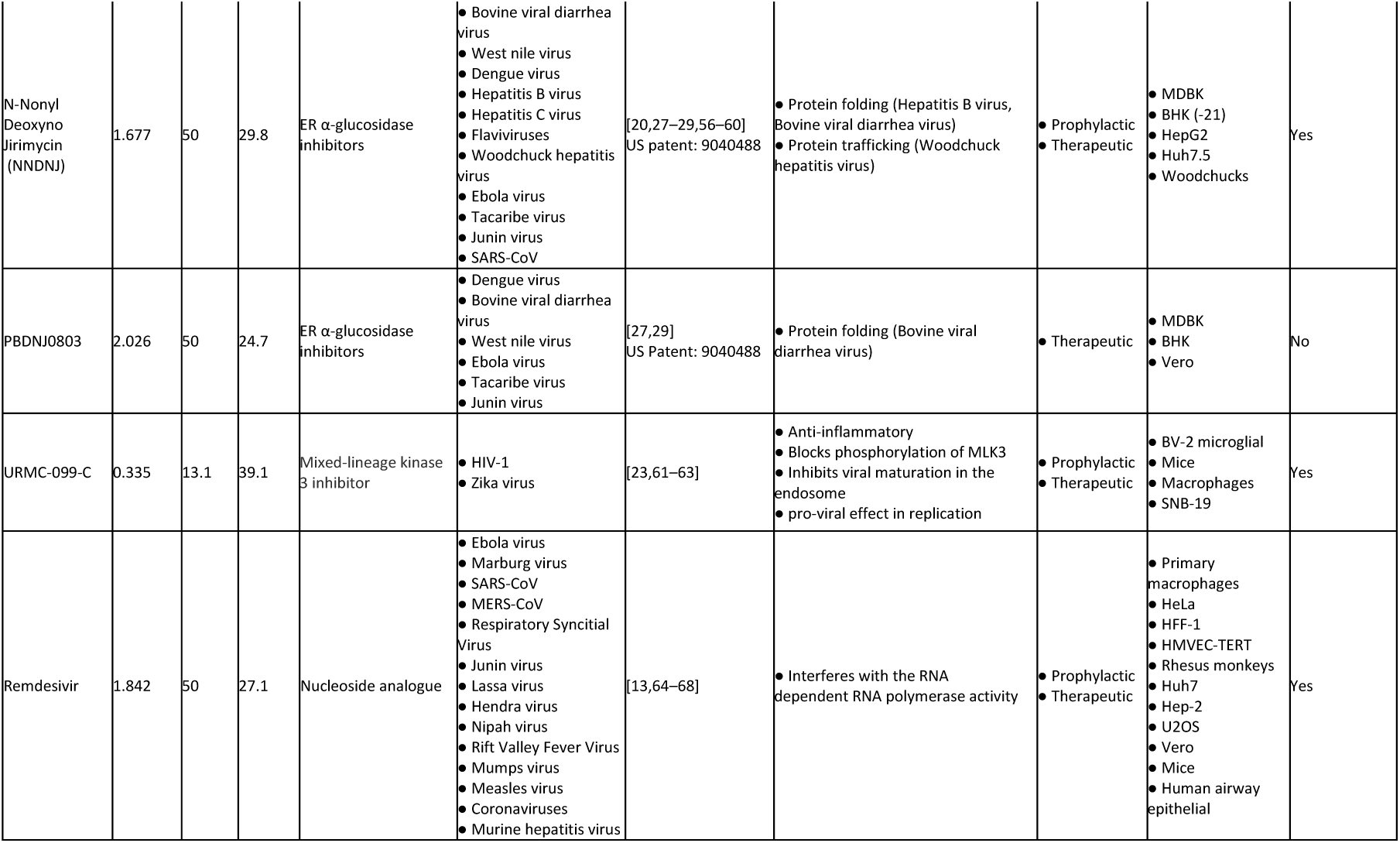
The IC_50_ and CC_50_ values of the 5 compounds that showed inhibition of SARS-CoV-2 during the compound dilution series, including references to the mode of action of these compounds against other viruses. Remdesivir is included as a reference drug control

| Name | IC<sub>50</sub> | CC<sub>50</sub> | SI | Class | Tested against other viruses | Literature reference | Mode of action | Administration | Tested on: | Bioavailable |
| --- | --- | --- | --- | --- | --- | --- | --- | --- | --- | --- |
| Retro-2.1 | 0.4909 | 50 | 101.9 | Retrograde transport inhibitor | • Enterovirus,<br>• Herpes simplex virus<br>• Filoviruses<br>• Vaccina virus<br>• Polyomavirus | [19,31–34] | • Modulate intracellular vesicle transport (Enterovirus 71),<br>• Blocks entry (Herpes Simplex Virus 2)<br>• Blocks entry that follows by glycoprotein proteolysis (Filoviruses)<br>• Localization towards the ER (Polyomavirus)<br>• endosome-to-Golgi apparatus trafficking (Vaccina virus)<br>• induce cellular STX5 displacement from the Golgi upon treatment, leading to an inhibition of retrograde transport (Cytomegalovirus) | • Prophylactic<br>• Therapeutic | • Vero<br>• HeLa<br>• 293S<br>• Mice | No |
| Chloroquine | 1.03 | 50 | 48.5 | Autophagic proteolysis inhibitor<br>Endosomal acidification inhibitor | • Herpes simplex virus<br>• Picornaviruses<br>• Poliovirus<br>• SARS-CoV<br>• HIV-1<br>• Hepatitis A virus<br>• Hepatitis B virus<br>• Hepatitis C virus<br>• Influenza A virus<br>• Influenza B virus<br>• Bovine viral diarrhea virus<br>• 229E-CoV<br>• OC43-CoV<br>• Chikungunya virus<br>• Dengue virus<br>• Crimean-Congo Hemorrhagic fever<br>• Ebola virus<br>• Nipah virus<br>• Hendra virus<br>• Lassa virus<br>• Rabies virus | [21,22,35–55] | • Raise endosomal pH (Picornavirus)<br>• Terminal glycosylation of the ACE2-receptor (SARS-CoV)<br>• Alter the glycosylation of the glycoprotein (HIV-1)<br>• Interference with endosomal acidification (Hepatitis C virus)<br>• Endosomal acidification (BVDV)<br>• P38 MAPK and ERK activation (229E) | • Prophylactic<br>• Therapeutic | • VeroE6<br>• Vero 76<br>• L132<br>• MDBK<br>• Huh7<br>• NS20 murine neuroblastoma cells<br>• B-SC-1<br>• A549<br>• Humans<br>• U937<br>• MRC5<br>• Guinea pigs | Yes |
| N-Nonyl<br>Deoxyno<br>Jirimycin<br>(NNDNJ) | 1.677 | 50 | 29.8 | ER α-glucosidase<br>inhibitors | • Bovine viral diarrhea virus<br>• West Nile virus<br>• Dengue virus<br>• Hepatitis B virus<br>• Hepatitis C virus<br>• Flaviviruses<br>• Woodchuck hepatitis virus<br>• Ebola virus<br>• Tacaribe virus<br>• Junin virus<br>• SARS-CoV | [20,27–29,56–60]<br>US patent: 9040488 | • Protein folding (Hepatitis B virus, Bovine viral diarrhea virus)<br>• Protein trafficking (Woodchuck hepatitis virus) | • Prophylactic<br>• Therapeutic | • MDBK<br>• BHK (-21)<br>• HepG2<br>• Huh7.5<br>• Woodchucks | Yes |
| PBDNJ0803 | 2.026 | 50 | 24.7 | ER α-glucosidase<br>inhibitors | • Dengue virus<br>• Bovine viral diarrhea virus<br>• West Nile virus<br>• Ebola virus<br>• Tacaribe virus<br>• Junin virus | [27,29]<br>US Patent: 9040488 | • Protein folding (Bovine viral diarrhea virus) | • Therapeutic | • MDBK<br>• BHK<br>• Vero | No |
| URMC-099-C | 0.335 | 13.1 | 39.1 | Mixed-lineage kinase<br>3 inhibitor | • HIV-1<br>• Zika virus | [23,61–63] | • Anti-inflammatory<br>• Blocks phosphorylation of MLK3<br>• Inhibits viral maturation in the endosome<br>• pro-viral effect in replication | • Prophylactic<br>• Therapeutic | • BV-2 microglial<br>• Mice<br>• Macrophages<br>• SNB-19 | Yes |
| Remdesivir | 1.842 | 50 | 27.1 | Nucleoside analogue | • Ebola virus<br>• Marburg virus<br>• SARS-CoV<br>• MERS-CoV<br>• Respiratory Syncytial Virus<br>• Junin virus<br>• Lassa virus<br>• Hendra virus<br>• Nipah virus<br>• Rift Valley Fever Virus<br>• Mumps virus<br>• Measles virus<br>• Coronaviruses<br>• Murine hepatitis virus | [13,64–68] | • Interferes with the RNA dependent RNA polymerase activity | • Prophylactic<br>• Therapeutic | • Primary macrophages<br>• HeLa<br>• HFF-1<br>• HMVEC-TERT<br>• Rhesus monkeys<br>• Huh7<br>• Hep-2<br>• U2OS<br>• Vero<br>• Mice<br>• Human airway epithelial | Yes |

**Figure 2:**
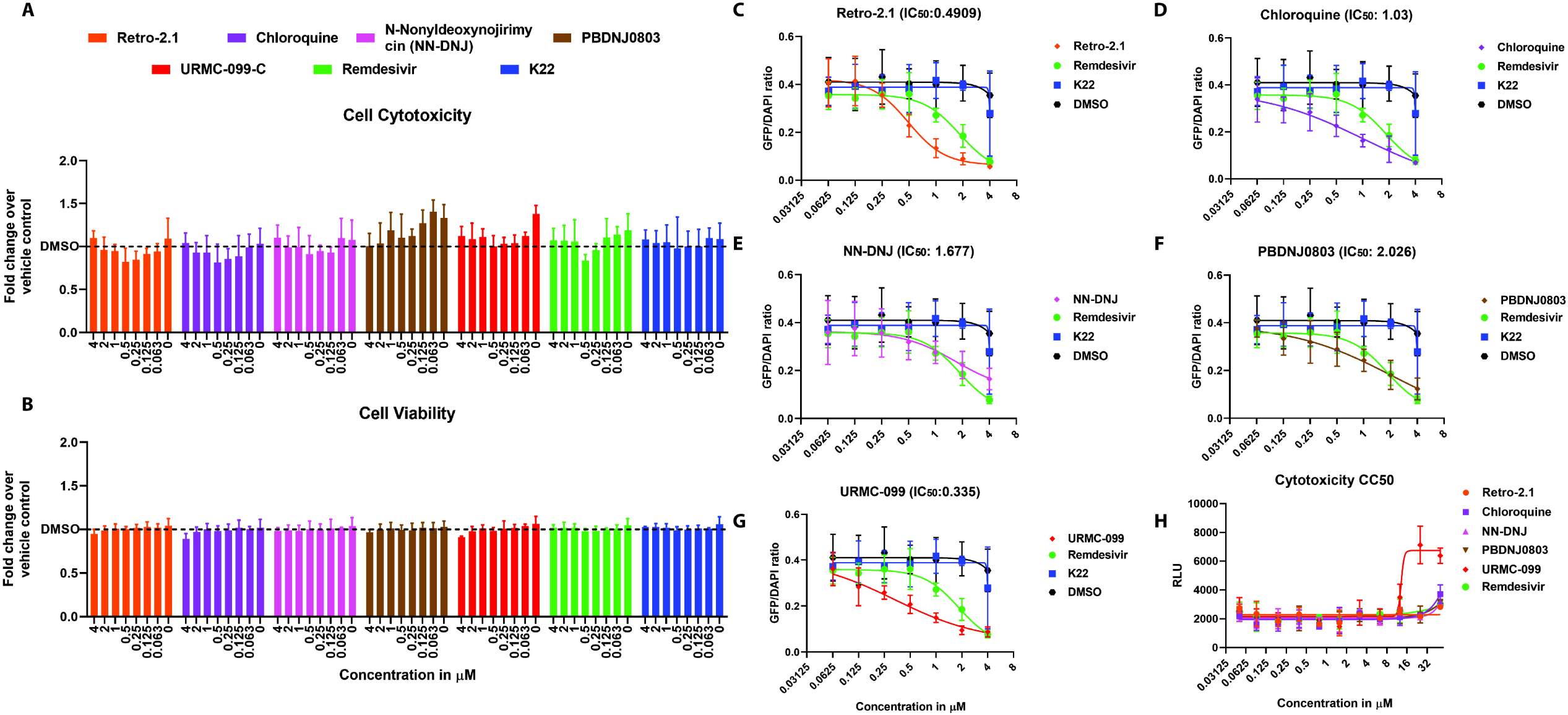
IC_50_ determination of the five compound hits that showed inhibition against SARS-CoV-2. The cell cytotoxicity **(A)** and cell viability **(B)** were assessed from each compound dilution of the five hits after 24 hours of incubation on Vero-E6 cells at 37°C with a 5% CO_2_-humidity. The cells were pretreated for 2 hours with the compound dilution prior to infection. To determine the inhibition of the five compounds, the measured total fluorescence signal intensity of GFP (infected cells) per well was divided by the total fluorescence signal intensity of DAPI (presence of cells) per well, indicated as the GFP/DAPI ratio. The GFP/DAPI ratio of each compound dilution of Retro-2.1 (**C)**, Chloroquine **(D)**, *n*-Nonyldeoxynojirimycin (NN-DNJ) **(E)**, PBDNJ0803 **(F)** and URMC-099 **(G**) are compared to Remdesivir, K22 and DMSO. Results are displayed as means and SD of three individual experiments. To control for cytotoxicity, the half maximum cytotoxicity concentration 50% (CC_50_) per compound were determined **(H)**. Results of the CC_50_ are shown as means and SD from two individual experiments.

In parallel to the efficacy, we determined the half-maximum cytotoxicity concentration (CC_50_) of each respective compound at a concentration range from 0.04 µM to 50 µM using uninfected Vero-E6 cells. This demonstrated that the previously tested compounds were all well tolerated at 2 or 4-fold higher concentrations than the initial screening concentration of 1 μM **(Figure 2H, table 2)**. At concentrations up to 50 µM, only URMC-099-C displayed moderate cell cytotoxicity, resulting in a CC_50_ of 13.1 µM, while all other compounds had a CC_50_ above 50 µM **(Table 2)**. Further, the resulting selectivity indexes (SI) showed that Retro-2.1 (SI = 101.9) is the most effective and less cytotoxic inhibitor, followed by chloroquine (SI = 48.5), URMC-099-C (SI = 39.1), whereas NN-DNJ (SI = 29.82), and PDNJ0803 (SI = 24.68) had comparable SI to that of Remdesivir (SI = 27.14) **(Table 2)**. Combined these results demonstrate that Retro-2.1 is the most potent antiviral candidate that should be further evaluated on its mode of action, efficacy and safety in pre-clinical models such as human airway epithelial cultures, as well as appropriate *in vivo* models for SARS-CoV-2.

## Discussion

In this study we demonstrate that a conservative *in vitro* screening approach of 400 compounds from the PRB resulted in the identification of five compounds with potent antiviral activity against SARS-CoV-2. This included the anti-malaria drug chloroquine and antibacterial Retro-2.1, as well as antiviral compounds NN-DNJ, PDNJ0803, and URMC-099-C. Antiviral efficacy testing revealed that Chloroquine, NN-DNJ, PDNJ0803, and URMC-099-C, all have a comparable characteristics to that of the reference compound Remdesivir, while the antimicrobial compound Retro-2.1 displayed almost a 4-fold higher selectivity index profile over that of Remdesivir in Vero-E6 cells. These results indicate that conservative *in vitro* screening approaches of preselected compounds can lead to the rapid identification of potent antiviral candidates targeting SARS-CoV-2. We and others have previously screened different compound libraries against a variety of coronaviruses in a range of 10 to 50 µM that identified several compounds with antiviral activity against SARS-CoV and MERS-CoV. [12,17,18]. However, in the current study we employed a stringent survival screening approach to expedite the identification of candidate compounds effective against SARS-CoV-2. This comes with the expense that compounds effective at higher molar concentrations are missed.

We used the nucleoside analogue Remdesivir as a reference compound against SARS-CoV-2 and observed an IC_50_ of 1.842 µM and a CC_50_ of >50 µM on Vero-E6 cells, in agreement with previous studies describing the inhibitory effect of Remdesivir against SARS-CoV-2 [8]. In line with this, we also identified Chloroquine as a potent antiviral compound against SARS-CoV-2, which has also been shown to effectively inhibit SARS-CoV-2 *in vitro* [8]. In contrast, K22 neither inhibited SARS-CoV nor SARS-CoV-2 **(Figure 1)** replication at low concentrations [12]. This exemplifies the robust reproducibility of the antiviral screen presented here and its consistency with previously published reports. Further, it establishes Remdesivir and chloroquine as reliable reference compounds during antiviral efficacy assessments of novel compounds against SARS-CoV-2 and other coronaviruses.

The five candidate compounds identified here, namely chloroquine, NN-DNJ, PDNJ0803, URMC-099-C, and Retro-2.1 have previously also been demonstrated to inhibit the replication of other RNA-viruses such as *Filoviruses, Flaviviruses* and *Picornaviruses* [19–21]. Although we did not evaluate the mode of action of the identified compounds, it has been described that Chloroquine interferes with the endosomal viral entry and release pathways of coronaviruses, including SARS-CoV-2, whereas the immunomodulatory compound URMC-099-C might influence important Mitogen-activated protein kinases (MAPK) associated biological processes [8,22,23]. Interestingly, Retro-2.1 remodels the intracellular distribution of syntaxins, which consequently alters vesicular retrograde transport between endosomes and the Golgi apparatus [24,25]. Of note, we previous identified several syntaxins (*stxbp1, stxbp3, stx5a*, and *stx18)* in close proximity of the replication and transcription complexes during coronavirus replication [26]. Since coronaviruses strongly rely on cellular endomembranes and trafficking pathways for efficient replication, Retro-2.1 might directly target the Achilles heel of coronavirus replication. Further, NN-DNJ and its derivate PBDNJ0803 are categorized into the same class of inhibitors that target the cellular α-glucosidase I and II, which support the maturation, secretion and function of viral glycoproteins [27]. NN-DNJ and PBDNJ0803 interfere with the interactions between glycoproteins and the ER-chaperones Calnexin and Calreticulin, thereby disturbing the removal of terminal glucose residues from the *n*-glycans of glycoproteins [28]. Moreover, NN-DNJ has been described to effectively inhibit the replication of a wide range of RNA viruses such as Hepatitis C virus and Flaviviruses, as well as SARS-CoV [27–29]. Lastly, URMC-099-C has mainly been described as a suppressor of the inflammatory response and has been suggested for antiretroviral therapy [30]. Investigations deciphering its effect during the viral life cycle are limited, however, it is involved in the MAPK-pathway and is a positive regulator of the c-jun N-terminal kinase signaling pathway [23]. In summary, further evaluation on the mode of action and *in vivo* efficacy of each compound against SARS-CoV-2 and other coronaviruses is warranted to establish whether conserved host or viral features can be efficiently targeted by the selected antiviral compounds.

In summary, the stringent *in vitro* screening of 400 compounds identified several candidate compounds possessing potent antiviral properties against SARS-CoV-2 that vindicate further efficacy and safety testing in pre-clinical *in vitro* and *in vivo* models as novel intervention strategies against SARS-CoV-2 and other coronaviruses.

## Acknowledgements

We would gratefully thank the University of Bern for providing special authorization to conduct our research during the SARS-CoV-2 outbreak, and the Medicine for Malaria Ventures (MMV, mmv.org) and the Drugs for Neglected Diseases initiative (DNDi, dndi.org) for their support, for the design of the pandemic response box and supplying the compounds. We also would like to thank Gary Prescott of Biotek Instruments (Part of Agilent) for support, help and training with the image acquisition and analysis.

