## Supplemental Figure Legends for "Identification of five antiviral compounds from the Pandemic Response Box targeting SARS-CoV-2"

**Supplementary Figure 1: The cell survival (A), cell cytotoxicity (B) and cell viability (C) of compounds in the pandemic response box.** The survival, cytotoxicity and viability were measured after 48 hours of incubation on Vero-E6 cells at 37°C with 5% CO_2_-humidity. **(A)** The heat-maps show the position on the plates where survival was observed with a MOI of 0.01, which was arbitrarily scored from 0-2, with 0 showing no viable cells and 2 showing virus-induced CPE resistant cells. The cell cytotoxicity **(B)** and cell viability **(C)** of each specific compound were compared to the DMSO-vehicle control from that specific experiment and the fold change over vehicle control is displayed in the heat maps. Results are shown as mean of two individual experiments performed in two technical replicates.

**Supplementary figure 2: IC50 determination of eleven compound hits that showed partial inhibition against SARS-CoV-2.** The cell cytotoxicity **(A)** and cell viability **(B)** of the remaining 11 compound hits that showed partial inhibition during the compound dilution series were analyzed after 24 hours of incubation on Vero-E6 cells at 37°C with a 5% CO_2_-humidity. The cells were pretreated for 2 hours with the compound dilution prior to infection. To determine the efficiency of the five compounds, the measured total fluorescence signal intensity of GFP (infected cells) per well was divided by the total fluorescence signal intensity of DAPI (presence of cells) per well, indicated as the GFP/DAPI ratio. The GFP/DAPI ratio of each compound dosage of Oteseconazole **(C)**, Itraconazole **(D)**, AC1MTT7T **(E)**, Sulbactam **(F)**, Decylphosphinate **(G)**, MMV1578576 **(H)**, NA for Racemic **(I)**, Lomefloxacin **(J)**, MMV1593544 **(K)**, Noscapine **(L)** and AZD0156 **(M)** are compared to Remdesivir, K22 and DMSO. Results are displayed as means and SD of three individual experiments.
