## Supplementary figures and images for "Identification of five antiviral compounds from the Pandemic Response Box targeting SARS-CoV-2"

### Supplemental Figure 1

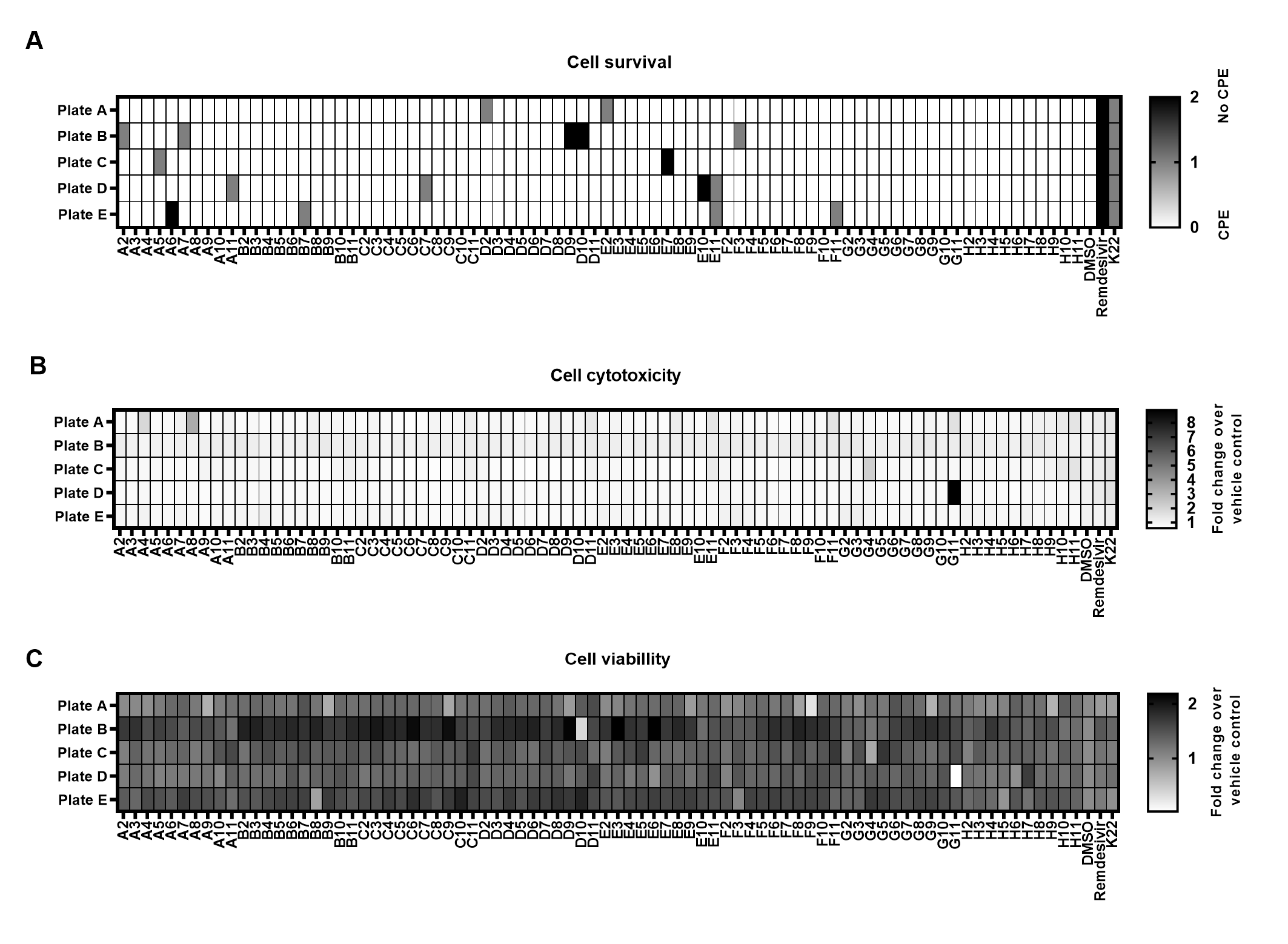

### Supplemental Figure 2

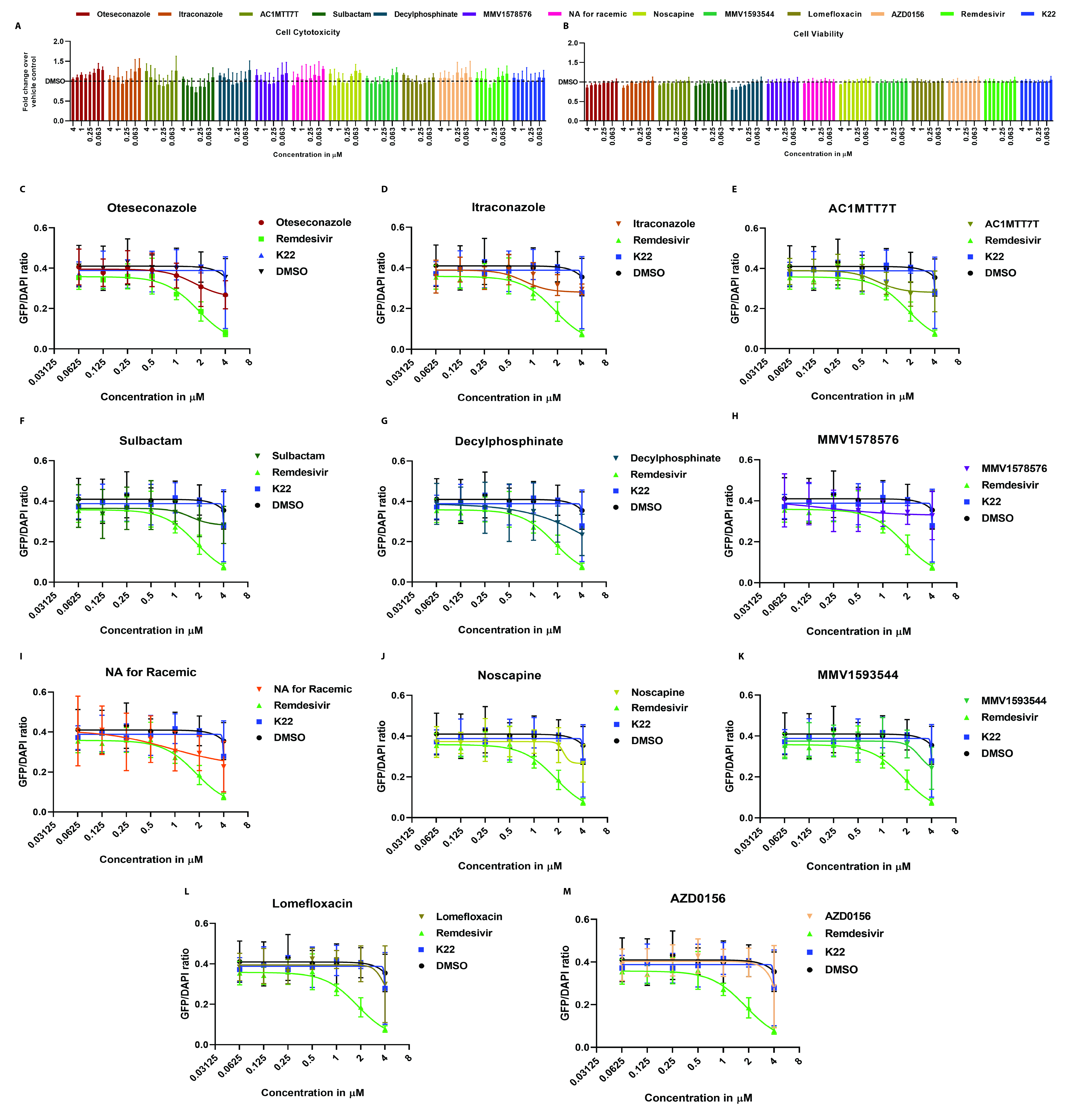
